# *In vitro* induced floral reversion in switchgrass (*Panicum Virgatum* L.)

**DOI:** 10.1101/871152

**Authors:** Yongfeng Wang, Kunliang Xie, Fengli Sun, Chao Zhang, Shudong Liu, Yajun Xi

**Author notes:** These authors contributed to the work equllly and should be regarded as co-first authors (Yongfeng Wang, Kunliang Xie). **Corresponding author:** Yajun Xi.

## Abstract

Switchgrass (*Panicum Virgatum* L.) is a warm-season perennial grass native to North America, it was used as forage and vegetative filter strips in early days, and have developed into a bioenergy crop in recent years. In this study, we found that the switchgrass cultivar ‘Alamo’ at elongation stage 4 have developed inflorescences about 1 cm in length, and *in vitro* incubation of the shoot apexes harboring inflorescences on Murashige and Skoog’s basal medium supplemented with 3 mg/L 6-benzylaminopurine generated multiple shoot clumps. Anatomical study showed that some of the regenerated shoots originated from axillary buds on the explants, some of them originated from adventurous buds and some of them originated from young florets. Further study of shoots originated from young florets found that the floral organs degenerated or developed into leaf-like organs, and the flower terminal transformed into a vegetative shoot apical meristem, that’s to say these shoots arise from flower reversion. *In vitro* induction of floral reversion provided a novel protocol to manipulate flower development in switchgrass, which might contribute a fundamental for flower development study in switchgrass and other plants.

## 1 Introduction

Flowering is an important process in plant life, through which hereditary material was transferred to offspring over sexual reproduction (Scutt & Vandenbussche, 2014). In general, plant species old enough carry out an irreversible one-way progression: blossom, pollinate, fertilize, and produce seeds (Irish 2010). However, some florally evoked plants may also reverse developed into an earlier phase or the vegetative phase if the photoperiod (Zhao et al., 2002; McCullough et al., 2010; Washburn & Thomas, 2000) or temperature (Day et al., 1994; Moncur, 1992) changed to a certain condition that was unfavorable for flowering, and it was termed flower reversion. Flower reversion has been described in many plants including both monocots and dicots (Battey & Lyndon, 1990; Tooke et al., 2005). According to the difference in reversion pattern, flower reversion was classified as flower reversion, inflorescence reversion and partial flowering (Battey & Lyndon, 1990).

Many factors have been reported influencing flower reversion, they were summarized in reviews of Battey and Lyndon (1990) and Tooke et al. (2005), and more recent reports are presented in this article as an up-date in table 1. Among factors influencing flower reversion, photoperiod and temperature were best described. For example, *Impatiens balsamina* is a short-day plant, transferring plants to continuous long-day after a treatment of 5 short days finally raise up to flower reversion (Tooke et al., 2005), *Arabis alpine* is a plant that requires 18-24 weeks of low temperature to be able to flower normally, with insufficient low temperature treatment, the florets on the inflorescence branches will convert to vegetative growth (Lazaro et al., 2018). Besides, the air humidity was also considered a factor influencing flower reversion. In *Whytockia bijieensis*, low temperature and low humidity in the night is the inductive condition of flower reversion, if the humidity is higher than 75%, the plant will not reverse developed (Wang, 2001). Plant hormones is another factor affecting flower development. In *I. balsamina*, insufficient short day treatment lead to flower reversion, however, exogenous application of gibberellin will suppress the transition (Nanda et al., 1967).

**Table 1.** Plant flower reversion and reversion feature in rencently publications

| Species | Flowering conditions | Reversion conditions | Characteristics of reverted plants | Reference |
| --- | --- | --- | --- | --- |
| *Dimocarpus longan* | Low temperature in winter | Warm temperature and high humidity | The inflorescence stopped reproductive growth and developed leaves, some of the inflorescence converted to vegetative shoots | (Chen et al., 2009) |
| *Arabidopsis suecica* | Long-day | Short-day | Flowers of this allotetraploid partially reversed in long days, and the frequency is greatly increased under short day condition | (McCullough et al., 2010) |
| *Oryza sativa* | Short-day | double mutation of *dep* and *afo* | In the double mutant of *dep* and *afo*, florets finally developed into shoots | (Wang et al., 2010) |
| *Glycine max* | Short-day | Long-day | By transfer plants grown in short days condition to long days condition for 2-5 weeks, and then back into short days, the flowering time was delayed and vegetative growth was found in florets and inflorescences | (Jiang et al., 2011) |
| *Arabis alpina* | Vernalization | Insufficient vernalization | Insufficient vernalization leads to the reverse development of flower primordia into new branches | (Lazaro et al., 2018) |
| *Allium sativum* | Not flower under natural condition | Reverse under natural condition | The inflorescence developed bulblet under natural condition | (Winiarczyk et al., 2018) |
| *Luculia gratissima* | Short-day | Long-day | By transfer plants grown in short days condition for 2-25days to continuous long days, the flower buds form vegetative buds, the bracts and sepals turned to be leaf-like | (Wan Y et al., 2018) |
| *Panicum virgatum* | Short-day | Down-regulate expression of *SPL7* and *SPL8* | Down-regulate any one of *SPL7* and *SPL8* result in flowering time delay, while down-regulate both lead to flower reversion | (Gou et al., 2019) |

Switchgrass is a warm-season perennial C4 plant native to North America, it was traditionally used in animal feed and soil erosion prevention (Parrish et al., 2012; Parrish & Fike, 2005). Due to the advantage of broad adaptation, drought tolerance, and high production of biomass, it was selected as a potential bioenergy crop by the United States Department of Energy in the 1980s (Keshwani & Cheng, 2009; Parrish et al., 2012; Parrish & Fike, 2005; Sanderson et al., 2006). In this study, *in vitro* incubation of switchgrass immature inflorescence produced flower reversion, and it was further confirmed by anatomy and histology study. Flower reversion induction provided a novel protocol to manipulate flower development in switchgrass, it will benefit us in switchgrass flower development and flower meristem maintain study.

## 2 Materials and Methods

### 2.1 Switchgrass floral reversion induction

The switchgrass cultivar ‘Alamo’ that grew under field conditions in Yangling, Shaanxi, China, was used as plant material. Shoot apexes of tillers at E3 to E4 stage (Moore et al., 1991; Xi et al., 2009) were harvested as explants. They were surface sterilized with 70% ethanol (Ante, Anhui, China) for 1 min and then with 8% sodium hypochlorite (Guanghua, Guangdong, China, active chlorine ≥ 5.5%) for 3 min (Burris et al., 2009; Chai et al., 2012). After three times rinse with sterilized reverse osmosis water, the two ends of each explant were removed 0.5 cm in length. The sterilized explants were subsequently split longitudinally, each part was placed on shoot induction medium (Chai et al., 2012).

The incubation was conducted under photoperiod of 20 h light 4 h dark and constant temperature of 25 °C. Four weeks later, the explants were transferred on fresh shoot induction medium for another 30 d of subculture (Chai et al., 2012).

Shoot induction medium: MS basal medium (Murashige & Skoog 1962), supplemented with 3 mg·L^-1^ BAP (Sanland, Fujian, China), 7.5 g·L^-1^ agar, and 30 g·L^-1^ sucrose (Chai et al., 2012).

### 2.2 Anatomical study of shoots

Shoots emerged from explants were separate from shoot clumps, and then they were distinguished by the origin and morphology characters. Different kinds of shoots were observed and compared under a stereoscopic microscope (Nikon SMZ1500, Japan). After that, shoots were split longitudinally with a scalpel, and the radial section was examined under a stereoscopic microscope as well.

### 2.3 Histological study of shoots

Shoots distinguished by morphology characters were fixed with formalin, acetic acid, and alcohol (FAA, formalin:acetic acid:70% alcohol = 5:5:90) stationary liquid for 6 hours, then they were stained with Ehrlich’s hematoxylin (Avwioro, 2011) (Maikun, Shanghai, China) for 3 days and immersed in tap water for 1 hour for differentiation. Stained materials were further dehydrated with a gradient alcohol series (20 min each of 70%, 80%, and 90%, followed by 15 min of 100% for twice), cleared with xylene, and infiltrated with melted paraffin wax (Yang, 1986). Then, they were embedded in a paraffin block and cut into 10 μm slices using a paraffin microtome (JinHuaHuiYou HY-202A, China). Slices were dewaxed with xylene, post-mounted with Permount Mounting Medium (HuShi, Shanghai, China), and analyzed under an optical microscope (Chongqing UOP, UB203i, China).

### 2.4 The analysis of factors influencing switchgrass flower reversion

The operations of explants harvesting and sterilization were conducted as described above. The BAP concertation in shoot induction medium was altered to 0, 1, 2, 3, 4 and 5 mg·L^-1^ in the study of BAP influencing switchgrass flower reversion. The day length was altered to 10, 12, 14, 16, 18 and 20 hours in the study of photoperiod influencing switchgrass flower reversion. In the study of *in vitro* culture influencing switchgrass flower reversion, the plants was grown in field conditions. Tillers at E3 to E4 stage were selected and sprayed with 0, 1, 3, 10, 20 and 50 mg·L^-1^ BAP, the heading date and the morphology of flowers were observed. Ten replicates were used for analyses described above.

## 3 Results

### 3.1 Shoot clumps induced from switchgrass immature inflorescence

At the experimental site (Shaanxi province, China), the switchgrass cultivar ‘Alamo’ sprouting in late April and heading in middle August. In July, the tillers developed into E3 to E4 stage with 3 or 4 nodes visible (Fig. 1a). Anatomical analysis revealed that an immature inflorescence about 1 cm in length had developed at the shoot apex (Fig. 1b). Histological study showed that these immature inflorescences had developed floret meristems, while the pistil and stamens were still missing (Fig. 1b and 1c). According to the development process of rice inflorescence, it was at the IN6 to IN 7 stage (Ikeda et al. 2004).

**Fig. 1.**
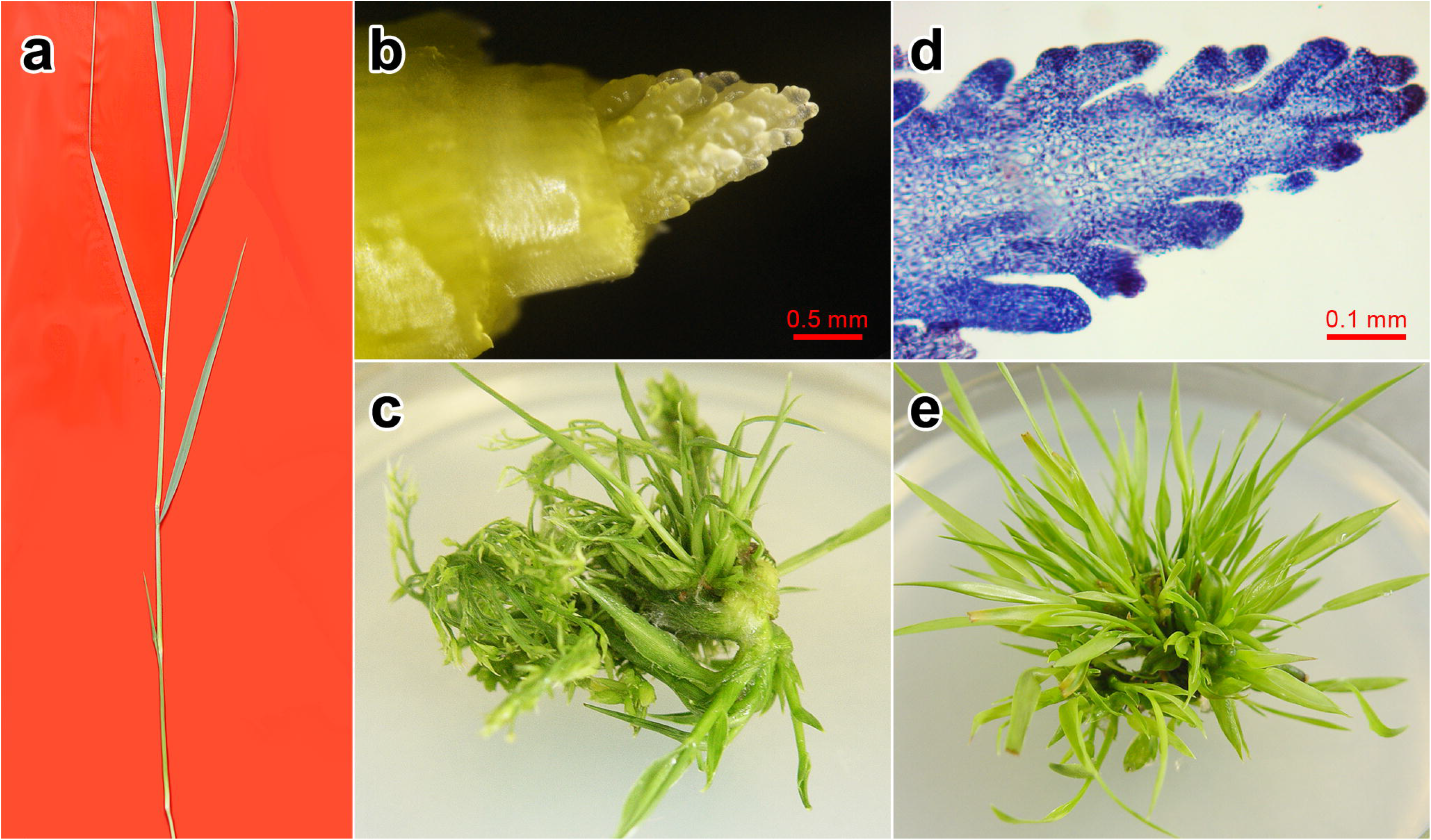
Induction of flower reversion in switchgrass at different stages. (a) Tiller at elongation stage 4; (b) Shoot apex excised from tiller; (c) Longitudinal section of shoot apex; (d) Shoot clumps emerged from explants after 30 day of incubation; (e) Shoot clumps emerged from explants after 60 day of incubation

After 30 days of incubation on shoot induction medium, explants developed into shoot clumps. The number of the shoots reach to 20 or more (Fig. 1d). As can be seen from figure 1d, shoots regenerated from different parts of explants were different in number and thickness. Shoots emerged from the base of the explants were stouter, while those emerged from apical side of the explants were thinner. However, shoots emerged from apical side were more dominant in quantity.

After another 30 days of subculture, the number of shoots emerged from an explant increased to 50 or more. And shoots emerged from different parts of the explants tend to be uniform in size (Fig. 1e).

### 3.2 Morphology analysis revealed three kinds of shoots with different origin

To investigate the origin of shoots arise from different parts of the explants, we detached the shoot clumps and compared the morphology characters of different kinds of shoots under a stereoscopic microscope, and a total of three kinds of shoots were found.

The first kind of shoots were speculated originated from the axillary buds, the evidence include: (1) This kind of shoots emerged from the base of the explants, where possessing two or three nodes with shortened internodes as described in other forage grasses (Fig.1b, Moore et al. 1991), and on the explants, axillary buds were also observed(Fig. 2a). (2) These shoots have a distinctive main stem and it was bigger in size compared with other shoots (Fig. 1d, Fig. 2b, c), which indicated that the main stem formed first while the other shoots formed relatively later. (3) These shoots were connected with the main stem directly and arranged symmetrically, which indicated that these shoots formed orderly along with the development of the main stem, which is similarly to the formation of axillary buds on the switchgrass tiller. To simplify these shoots were named as “AX shoots” in the following description.

**Fig. 2.**
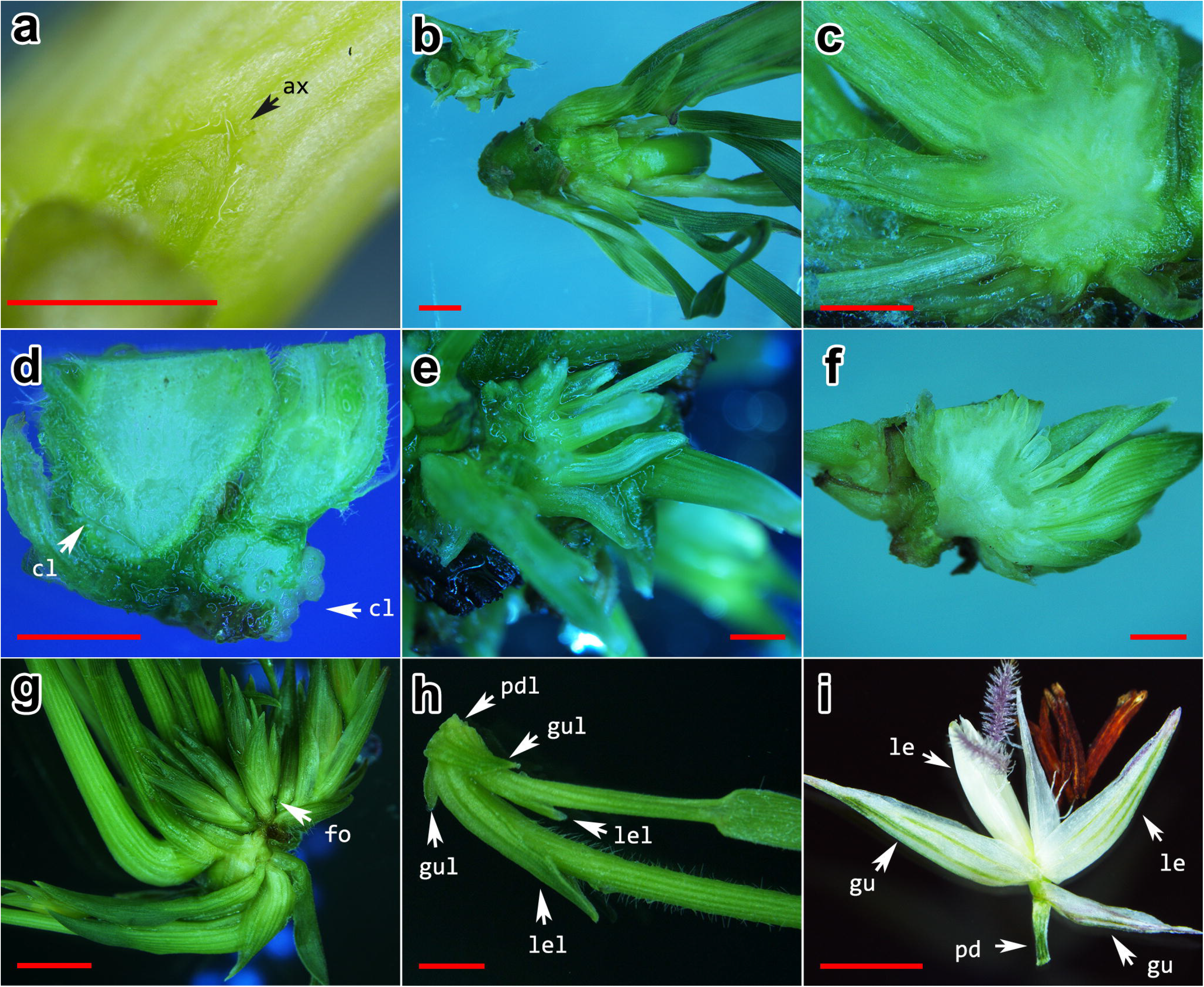
Morphology features of shoots from switchgrass shoot apexes. (a) Axillary bud on explants; (b) Shoots emerged from axillary buds; (c) Longitudinal section of shoots emerged from axillary buds; (d) Callus emerged from nodes; (e) Shoots emerged from adventitious buds; (f) Longitudinal section of shoots emerged from adventitious buds; (g) Shoots emerged from flower buds; (h) Shoots separated from shoot clump emerged from flower buds; (i) A normal floret of switchgrass. Bars = 2 mm Cl: callus; fo: floret; pd: pedicel; pdl: pedicel-like; gu: glume; gul: glume-like; le: lemma; lel: lemma-like

Another kind of shoots were speculated arise from adventitious buds, the evidence include: (1) Explants used in this experiment were shoot apexes harvested at the E3-E4 stage, they were less differentiated organs, and callus was observed on explants accompanying with the shoot clumps formation (Fig. 2d). (2) This kind of shoots grown out from the same position of the explants. As shown in figure 2e and 2f, about 10 shoots emerged from the node of explant, and they merged together at the base of the shoot clump, which indicated that they were arise from the callus induced from the node of the explants. To simplify these shoots were named as “AD shoots” in the following description.

There was another kind of shoots emerged from the explants, they were speculated originated from flower reversion, the evidence include: (1) Some floret like organs were observed accompanying with these shoots (Fig. 2g), and they were connected at the base. (2) These shoots arranged different from AX and AD shoots, they were connect at the base but didn’t merged like AD shoots, and they arranged dispersedly like florets on a spikelet (Fig. 2g). (3) There are several similarities between the third kind of shoots and the normal floret of switchgrass (Fig. 2h and 2i). Switchgrass spikelet was two flowered (Fig. 2i, Martinez-Reyna & Vogel, 1998), which contain a sterile floret with only three stamens and a fertile floret with three stamens and one pistil (Supplementary Fig. S1). In the regenerated shoots, we can also recognize two associated shoots grown out (Fig. 2h). At the base of these shoots, we can find a smallish and shortened tissue (Fig. 2h) which like the pedicel in a normal floret (Fig. 2i). There were two small leaves bent inward at the base of these shoots (Fig. 2h), they were similar to glumes of a floret (Fig. 2i). And at the bottom of each shoot, we also observed a bract-like leaf (Fig. 2h) which corresponding to the lemma in a floret (Fig. 2i). Morphological characters described above inferred that this kind of shoot might arise from the florets of switchgrass immature inflorescence, and to simplify it was named as “FL shoots”.

### 3.3 Histological analysis revealed FL shoots arise from flower reversion

Although we can confirmed that Fl shoots arise from floret, it was difficult to say if FL shoots initiated from switchgrass flower reversion or somatic embryos from florets due to the pluripotency of plant cells. For further understanding of the origin of FL shoots, here we analyzed the histological characteristics of FL shoots.

Figure 3a showed longitudinal section of two concomitant FL shoots. In which the residues of floral organs like glumes, lemmas and paleae were recognizable. From figure 2h and 3a, we can found that the concomitant FL shoots were different in size. According to the floral-like leaf arrangement on FL shoot and corresponding floral organs arrangement on floret showed in figure 2i, it was recognized that the stouter FL shoot was initiated from the second floret with both stamens and pistil while the other one was initiated from the first floret with only stamens (Martinez-Reyna & Vogel, 1998). In FL shoot initiated from the fertile floret (Fig. 3a, left), the inner whorls tissues stopped develop into floral organs. The stamens degenerated and shortened stamen-like tissues retained. At the terminal of the floret, pistil was replaced by a newly-formed tissue without style or stigma. While in FL shoot initiated from the sterile floret (Fig. 3a, right), the stamens were also degenerated. And at the terminal of the floret, which was degenerated in normal florets (Supplementary Fig. S1), cells resume growth and formed a bulge, it was speculated the growing point of the regenerated shoot. No callus or somatic embryos were observed in FL shoots.

**Fig. 3.**
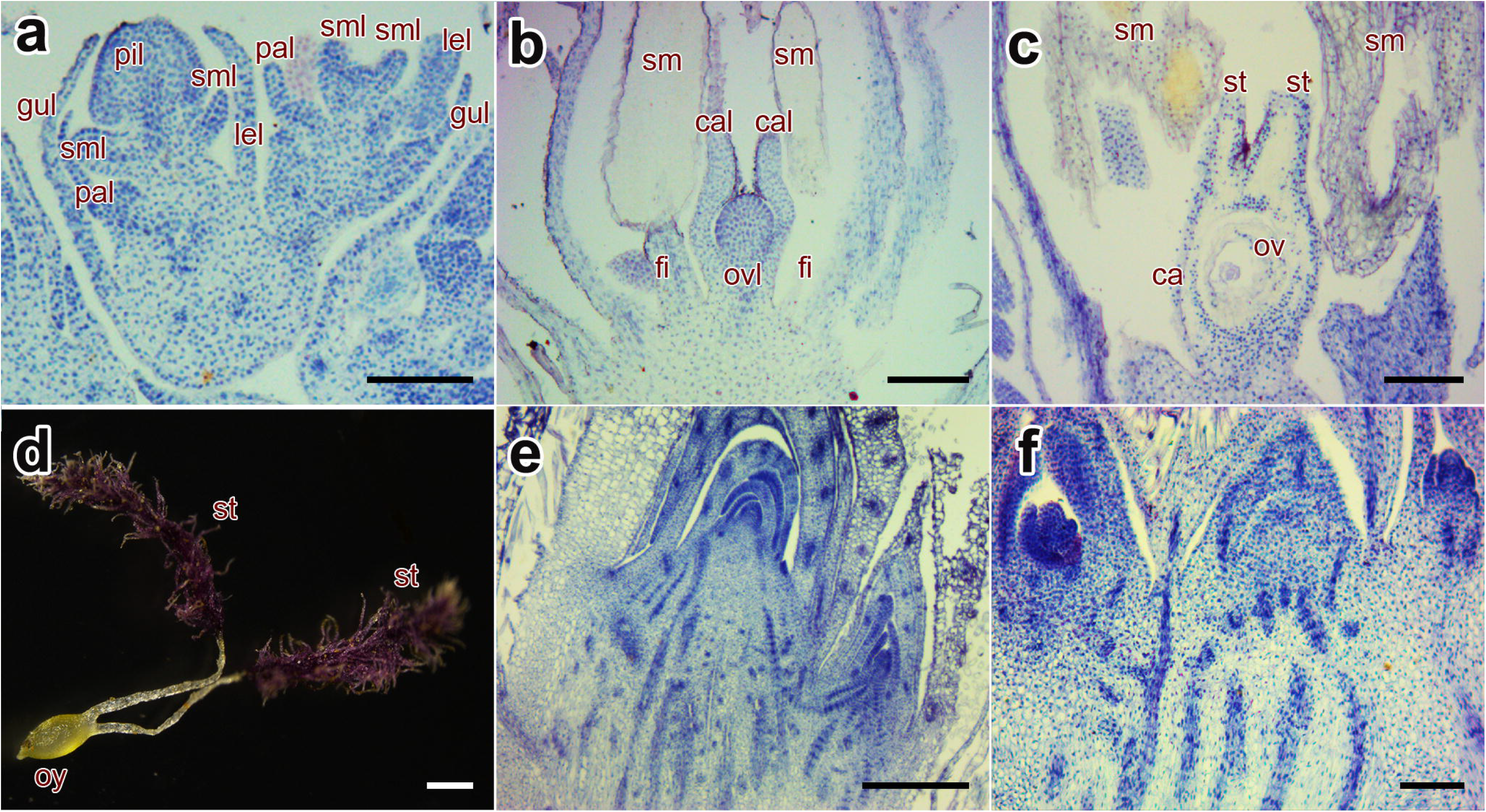
Histological features of shoots from switchgrass shoot apexes. (a) Longitudinal section of shoots emerged from flower buds; (b) Longitudinal section of half-reversed floret; (c) Longitudinal section of un-reversed floret; (d) A normal pistil of switchgrass; (e) Longitudinal section of shoots emerged from axillary buds; (f) Longitudinal section of shoots emerged from adventitious buds. In d and e, bar = 400μm, in other parts, bar = 100μm Ca: carpel; cal: carpel-like; gul: glume-like; pal: palea-like; lel: lemma-like; sml: stamen-like; pil: pistil-like; sm: stamen; ov: ovule; ovl: ovary-like; st: style; fi: filament; oy: ovary

In figure 3b and 3c, we compared the microstructure of florets adjacent to FL shoots (Fig. 2g) and a normal floret (Fig. 2i). Result showed that florets adjacent to FL shoots displayed partial reversion. In these florets, the filaments were obviously shortened, and at the inner whorl where pistils should be grown out, two leaf-like tissue were observed. According to the development of flower organ in rice (Supplementary Fig. S2, Ikeda et al., 2004), they might developed from the carpel primordium although they were not fused together like a normal ovary (Fig. 3c). At the center of the normally developed floret, we can find an ovary with two separately growing styles at the top and an ovule in the center (Fig. 3c and 3d), yet at the center of the half reversed floret, a globoid tissue with neither style nor ovule was formed (Fig. 3b). Since it located at the inner whorl of the carpel, we speculated the globoid tissue was developed from the ovule primordium. Callus and somatic embryos were also not found in the half reversed florets.

To validate that histological characteristics mentioned above were unique in FL buds and half reversed florets, we further analyzed the histological structure of AX and AD shoots. In AX shoot clumps, the growing points of apical bud and axillary buds were small, they were covered by layers of young leaves (Fig. 3e). This structure was the same as shoot apical meristem in monocot plants like rice and maize. While in AD shoot clumps, the shoots arranged side by side, and the shoot apical meristems were also covered by young leaves although they were less compact than that of AX shoots (Fig. 3f). No floral-shaped tissue or organ was observed in both AX and AD shoots.

Combine results mentioned above we can conclude that FL shoots arise from floret primordium, and they were initiated from growth pattern change of the floret primordium, which means they were developed from flower reversion rather than somatic embryo.

### 3.4 Factors influencing switchgrass flower reversion

Flowering is a complex phenomenon regulated by multiple factors, of which photoperiod, temperature, and growth regulators have been widely investigated and showed common effects in most plant species. In this study, the living condition of reverse developed young inflorescences changed in several aspects compared with the normal ones developed outdoor. To explore the importance of each factor, we further analyzed their effect on flower reversion independently.

#### High concentration of BAP facilitated switchgrass flower reversion

To explore the effect of BAP on switchgrass flower reversion induction, we analyzed the number of three kind of shoots under different concentration of BAP. Results showed that the number of AX shoot increased significantly when the level of BAP was increased from 0 to 3 mg.L^-1^, while it changed little when the BAP concentration increased from 3 to 5 mg.L^-1^. No AD shoot was induced when the BAP concentration was lower than 1 mg.L^-1^, however the number of AD shoot increased observably at higher BAP concentrations (*≥* 2 mg.L^-1^) and it reached a maximum at 4 mg.L^-1^ (Fig. 4a). FL shoot was not induced when the BAP concentration was lower than 1 mg.L^-1^. When the BAP level increased to 2 mg.L^-1^, a small number of FL shoots started to emerge. When it increased to 3 mg.L^-1^, the number of FL shoots increased significantly, the average number reach to 8.9 shoots per explant, and it changed little at higher BAP concentrations (4 and 5 mg.L^-1^).

**Fig. 4.**
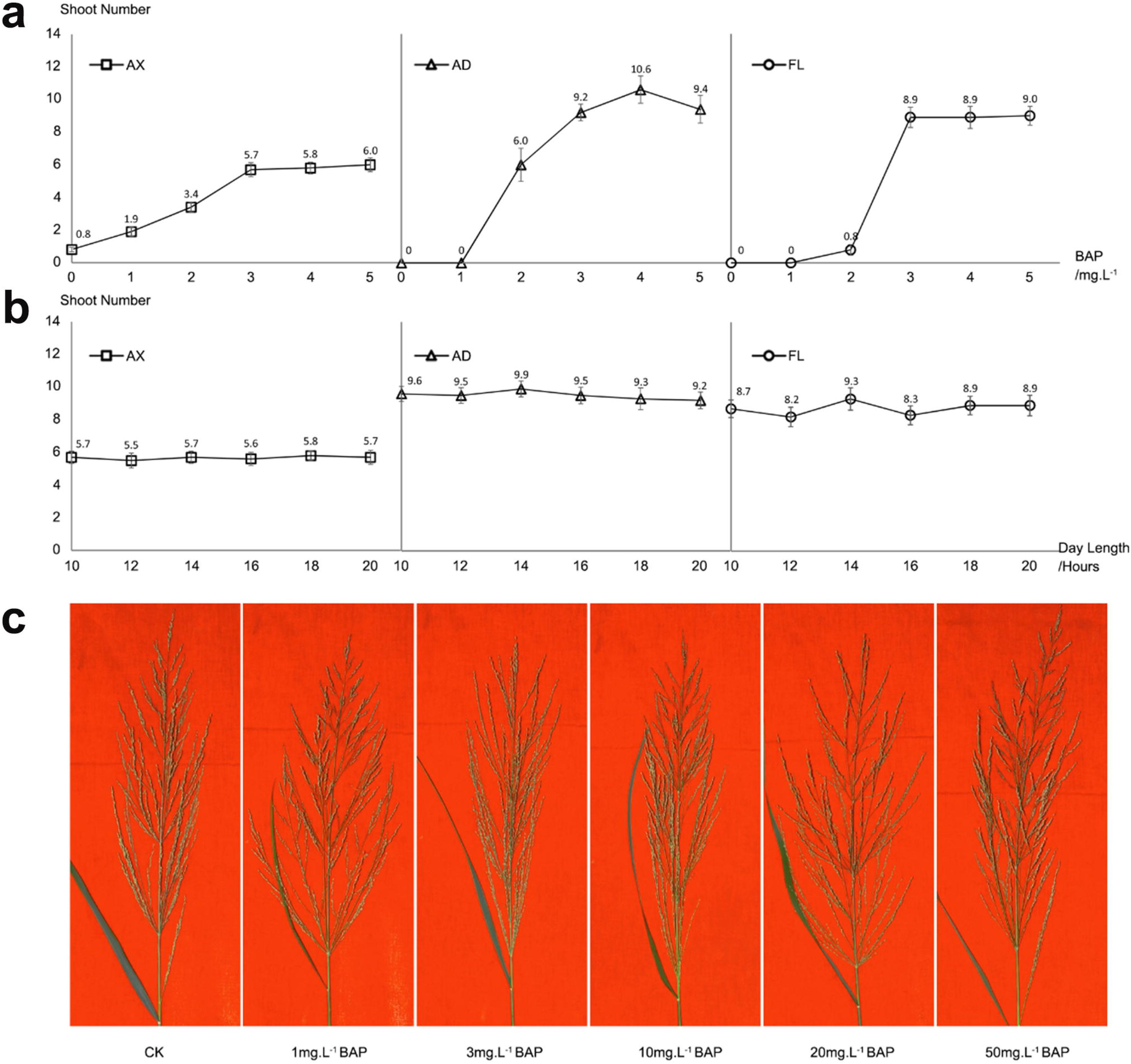
Analysis of factors influencing switchgrass flower reversion. (a) The effect of BAP on shoot number of three kinds of shoots; (b) The effect of day length on shoot number of three kinds of shoots; (c) The effect of BAP on switchgrass inflorescence development in plant

#### The photoperiod have little effect on switchgrass flower reversion

The day length used in this study was elongated to 20 hours compared with the filed condition (14 hours). To explore if the photoperiod played an important role in switchgrass flower reversion, we analyzed the induction of three kind of shoots under a set of day time from 10 hours to 20 hours. Results showed that the number of AX, AD and FL shoots changed a little under different photoperiods (Fig. 4b), which suggesting that the day length was less important in switchgrass flower reversion.

#### In vitro culture is required for switchgrass flower reversion

The living condition of explants in switchgrass flower reversion induction was different compared with the plants in filed condition. To explore if the *in vitro* environment was essential for switchgrass flower reversion, we analyzed the effect of BAP on switchgrass inflorescence development in filed condition. As shown in figure 4c, the development of switchgrass inflorescence was not effected by exogenously supplied BAP, and no flower reversion event was observed on switchgrass inflorescences under different BAP concentrations. Which suggesting that the *in vitro* environment was crucial for switchgrass flower reversion induction.

## 4 Discussion

Flower reversion is an abnormal development of floral organs, it was affected by various physiological and hormonal factors. In this study, we achieved flower reversion in switchgrass by incubating young inflorescences on shoot induction medium. Anatomical and histological study showed that shoots emerged from the explants originated in three ways: axillary buds, adventitious buds and floral buds. And shoots came from floral buds arise from the reverse development of flower primordia.

Flowering is a complex phenomenon regulated by multiple factors, of which photoperiod, temperature, and growth regulators have been widely investigated and showed common effects in most plant species. Over the past decades, many studies have been carried out to reveal the underlying mechanisms of flower initiation and floral organ determination (Kramer & Hall, 2005; Li et al., 2003; Purugganan et al., 1995), both environmental factors (e.g., photoperiod and temperature) and internal factors (e.g., cytokinin, auxin and jasmonic acid) have been demonstrated affect flower formation. In this study, explants incubated on MS medium without BAP or with low concentration or BAP could not generate flower reversion, which suggested that the BAP is a key factor promoting flower reversion. Whereas, it was reported that 50 μM BAP promote flowering in Arabidopsis (D’Aloia et al., 2011), this is probably because they used a transient treatment of 8 hours while a sustained treatment was used in this study. The number of three kinds of shoots were not influenced by the photoperiod, which indicated the photoperiod was not crucial in switchgrass flower reversion despite its important role in flower initiation (Esbroeck et al., 2003; Schwartz & Amasino, 2013). Moreover, the young inflorescences in plants were not reverse developed even if high concentration of BAP was supplemented, which indicated that the BAP was not sufficient for switchgrass flower reversion, and the rich organic inorganic and matters in the *in vitro* culture were also necessary factors in switchgrass flower reversion induction.

The panicles of switchgrass developed basipetally, which means florets near the bottom of the inflorescence are younger than those at the top (Martinez-Reyna & Vogel, 1998). In this study, we found that the recognized floral reversion mostly take place at the base of the immature inflorescence, which indicated that florets at early developmental stages were easier to reverse than those at a late stage. Moreover, in the half-reversed florets adjacent to FL shoots, it was found that the pistil showed reverse development while the stamen developed straightly, as it was well known that the formation of pistils occurs later than stamens and petals (Zik & Irish, 2003), we can deduced that floral reversion originated from the youngest tissues in a floret.

Floral reversion is an interesting phenomenon in plant development. However, most of the previous research concerning flower reversion focused on physiological factors influencing flower development, while few molecular biology investigations were conducted. The most likely limitation is the instability of *in vivo* induction of flower reversion described in previous reports (Battey & Lyndon, 1990; McCullough et al., 2010; Tooke et al., 2005). The *in vitro* induction of switchgrass flower reversion provide us a relatively stable protocol, and might contribute to reveal the underlying mechanisms of flower reversion and the maintenance of floral organs.

## Acknowledgments

The authors would like to thank Mr. Bu Bin for the guidance in paraffin sectioning and Dr. Quan Li for editing this manuscript. This work was supported by the National Natural Science Foundation of China (31171607), National Natural Science Foundation Committee.

---

**Figure S1.**
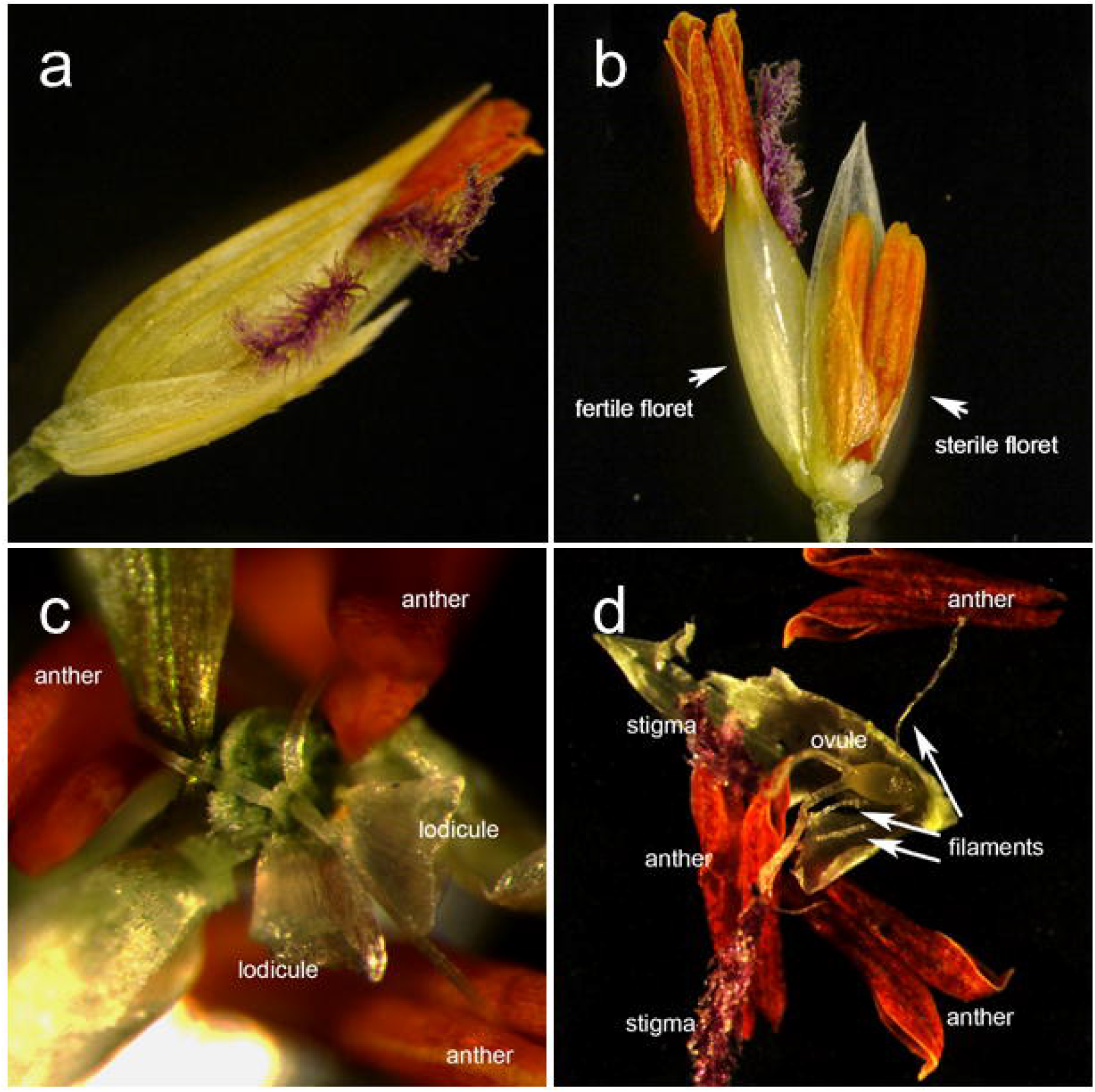
Switchgrass flower. (a). Overview of a switchgrass flower. (b). The fertile floret and sterile floret in switchgrass flower. (c). The Vertical view of sterile floret. (d). The side view of fertile floret.

**Figure S2.**
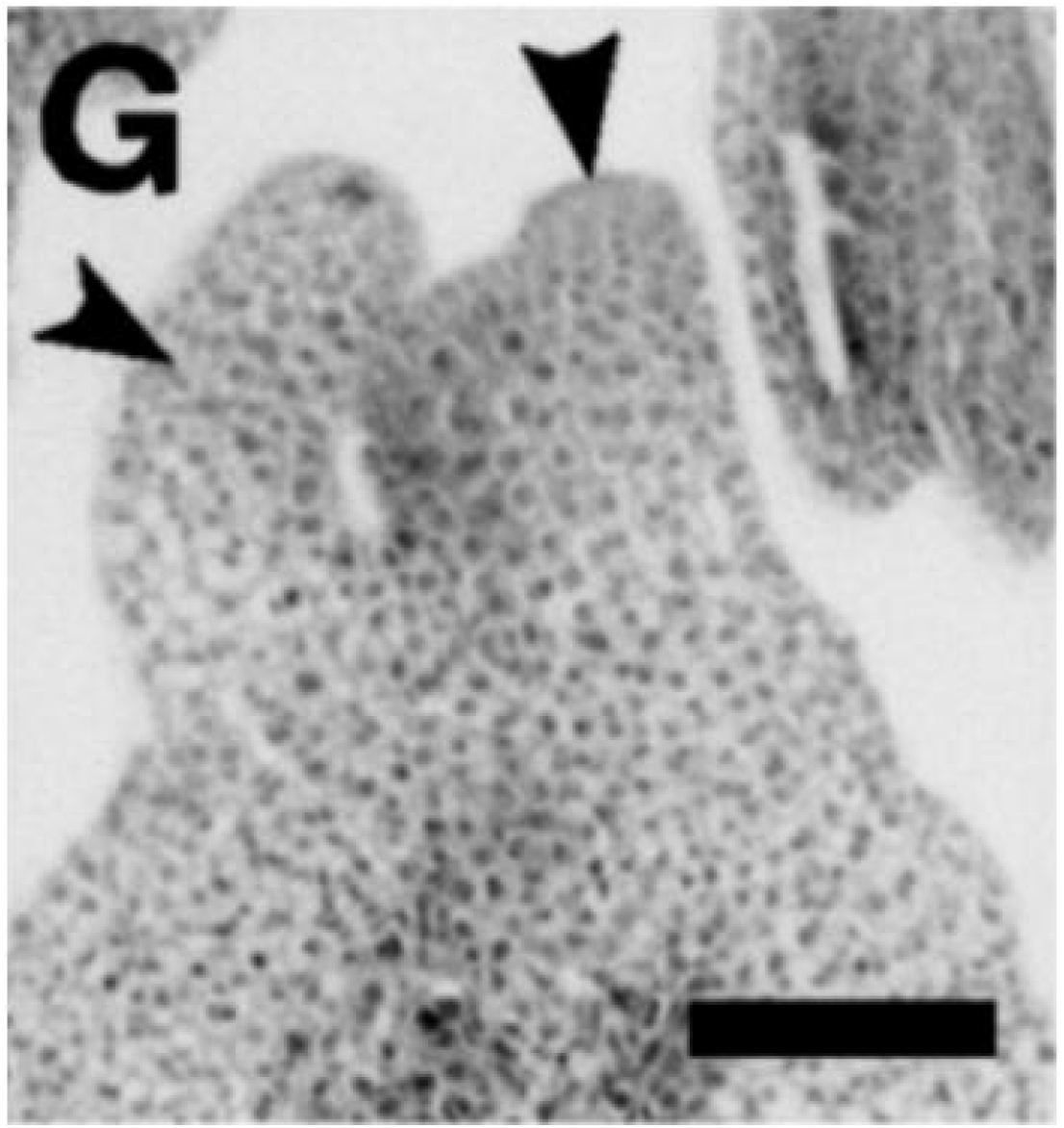
(G) Carpel primordium (arrowhead) formation of rice. A. Ikeda, K., Sunohara, H., and Nagato, Y. (2004). Developmental course of inflorescence and spikelet in rice. Breeding Sci 54(2), 147-156. doi: DOI 10.1270/jsbbs.54.147.

## REFERENCES

Avwioro, G. (2011) Histochemical Uses of Haematoxylin - A Review. Journal of Physics and Chemistry of Solids, 1, 24–34.

Battey, N.H. & Lyndon, R.F. (1990) Reversion of flowering. Botanical Review, 56, 162–189 doi:10.1007/BF02858534

Burris, J.N., Mann, D.G., Joyce, B.L. & Stewart. C.N. (2009) An improved tissue culture system for embryogenic callus production and plant regeneration in switchgrass (*Panicum virgatum* L.). BioEnergy Research, 2, 267–274 doi:10.1007/s12155-009-9048-8

Chai, G., Xu, K., Wang, Y, Sun, F., Song, L., Liu, S. & Xi, Y. (2012) Establishment of a highly efficient regeneration system of switchgrass artificial spike buds. Acta Prataculturae Sinica, 4, 98–104

Chen S, Liu H, Chen W, Xie D & Zheng S (2009) Proteomic analysis of differentially expressed proteins in longan flowering reversion buds. Scientia Horticulturae, 122, 275–280 doi:10.1016/j.scienta.2009.05.015

D’Aloia, M., Bonhomme, D., Bouché, F., Tamseddak, K., Ormenese, S., Torti, S., Coupland, G. & Perilleux, C. (2011) Cytokinin promotes flowering of Arabidopsis via transcriptional activation of the FT paralogue TSF. The Plant Journal, 6, 972–979 doi:10.1111/j.1365-313X.2011.04482.x

Day, J., Loveys, B. & Aspinall, D. (1994) Environmental control of flowering of *Boronia megastigma* (Rutaceae) and *Hypocalymma angustifolium* (Myrtaceae). Australian journal of botany, 42, 219–229 doi:10.1071/BT9940219

Esbroeck, G.A., Husseyab, M.A. & Sanderson, M.A. (2003) Variation between Alamo and Cave-in-Rock Switchgrass in Response to Photoperiod Extension. Crop Science, 43, 639–643

Gou, J., Tang, C., Chen, N., Wang, H., Debnath, S., Sun, L., Flanagan, A., Tang, Y., Jiang, Q., Allen, R.D. & Wang Z. (2019) SPL7 and SPL8 represent a novel flowering regulation mechanism in switchgrass. New Phytologist, 222, 1610–1623 doi:10.1111/nph.15712

Ikeda, K., Sunohara, H. & Nagato, Y. (2004) Developmental course of inflorescence and spikelet in rice. Breeding Science, 54, 147–156 doi:10.1270/jsbbs.54.147

Irish, VF. (2010) The flowering of Arabidopsis flower development. The Plant journal, 61, 1014–1028 doi:10.1111/j.1365-313X.2009.04065.x

Jiang, Y., Wu, C., Zhang, L., Hu, P., Hou, W., Zu, W. & Han, T. (2011) Long-day effects on the terminal inflorescence development of a photoperiod-sensitive soybean *[Glycine max* (L.) Merr.] variety. The Plant journal, 180, 504–510 doi:10.1016/j.plantsci.2010.11.006

Keshwani, D.R. & Cheng, J.J. (2009) Switchgrass for bioethanol and other value-added applications: A review. Bioresource Technology, 100, 1515–1523 doi:10.1016/j.biortech.2008.09.035

Kramer, E.M. & Hall, J.C. (2005) Evolutionary dynamics of genes controlling floral development. Current Opinion in Plant Biology, 8, 13–18 doi:10.1016/j.pbi.2004.09.019

Lazaro, A., Obeng-Hinneh, E. & Albani, M.C. (2018) Extended Vernalization Regulates Inflorescence Fate in *Arabis alpina* by Stably Silencing PERPETUAL FLOWERING1. Plant physiology, 176, 2819–2833 doi:10.1104/pp.17.01754

Li, G.S., Meng, Z., Kong, H.Z., Chen, Z.D. & Lu, A.M. (2003) ABC model and floral evolution. Chinese Science Bulletin, 48, 2651–2657 doi:10.1007/BF02901752

Martinez-Reyna, J.M. & Vogel, K.P. (1998) Controlled hybridization technique for switchgrass. Crop Science, 38, 876–878 doi:10.2135/cropsci1998.0011183X003800030042x

McCullough, E., Wright, K.M., Alvarez, A., Clark, CP., Rickoll, W.L. & Madlung, A. (2010) Photoperiod-dependent floral reversion in the natural allopolyploid Arabidopsis suecica. New Phytologist, 186, 239–250 doi:10.1111/j.1469-8137.2009.03141.x

Moncur, M. (1992) Effect of low temperature on floral induction of Eucalyptus lansdowneana F. Muell. & J. Brown subsp. Lansdowneana. Australian Journal of Botany, 40, 157–167 doi:10.1071/BT9920157

Moore, K.J., Moser, L.E., Vogel, K.P., Waller, S.S., Johnson, B.E. & Pedersen, J.F. (1991) Describing and quantifying growth stages of perennial forage grasses. Agronomy Journal, 83, 1073–1077 doi:10.2135/cropsci1995.0011183X003500010007x

Murashige, T. & Skoog, F. (1962) A revised medium for rapid growth and bio assays with tobacco tissue cultures. Physiologia plantarum, 15, 473–497 doi:10.1111/j.1399-3054.1962.tb08052.x

Nanda, K.K., Krishnamoorthy, H.N. & Anuradha, T.A. (1967) Floral induction by gibberellic acid in *Impatiens balsamina*, a qualitative short-day plant. Planta, 76, 367–370 doi:10.1007/BF00387542

Parrish, D.J., Casler, M.D. & Monti, A. (2012) The evolution of switchgrass as an energy crop. In: Monti A (ed) Switchgrass, A Valuable Biomass Crop for Energy. Green Energy and Technology. Springer-Verlag, London, pp 1–28. doi:10.1007/978-1-4471-2903-5_1

Parrish, D.J. & Fike, J.H. (2005) The biology and agronomy of switchgrass for biofuels. Critical Reviews in Plant Sciences, 24, 423–459 doi:10.1080/07352680500316433

Purugganan MD, Rounsley SD, Schmidt RJ & Yanofsky MF (1995) Molecular evolution of flower development: diversification of the plant MADS-box regulatory gene family. Genetics, 140, 345–356

Sanderson, M.A., Adler, P.R., Boateng, A.A., Casler, M.D. & Sarath, G. (2006) Switchgrass as a biofuels feedstock in the USA. Canadian Journal of Plant Science, 86, 1315–1325 doi:10.4141/P06-136

Schwartz, C. & Amasino, R. (2013) Nitrogen recycling and flowering time in perennial bioenergy crops. Frontiers in Plant Science, 4, 76 doi:10.3389/fpls.2013.00076

Scutt, C.P. & Vandenbussche, M. (2014) Current trends and future directions in flower development research. Annals of Botany, 114, 1399–1406 doi:10.1093/aob/mcu224

Tooke, F., Ordidge, M., Chiurugwi, T. & Battey, N. (2005) Mechanisms and function of flower and inflorescence reversion. Journal of Experimental Botany, 56, 2587–2599 doi: 10.1093/jxb/eri254

Wan, Y., Ma, H., Zhao, Z., Li, T., Liu, X., Liu, X. & Li, Z. (2018) Flowering response and anatomical study on process of flower bud differentiation for *Luculia gratissima* ‘Xiangfei’ under different photoperiods. Xibei Zhiwu Xuebao, 38, 1659–1666

Wang, Y. (2001) Reversion of floral development under adverse ecological conditions in *Whytockia bijieensis* (Gesneriaceae). Australian Journal of Botany, 49, 253–258 doi:10.1071/bt00002

Wang, K., Tang, D., Hong, L., Xu, W., Huang, J., Li, M., Gu, M., Xue, Y., Cheng, Z. (2010) DEP and AFO Regulate Reproductive Habit in Rice. PLoS Genetics, 6, e1000818 doi:10.1371/journal.pgen.1000818

Washburn, C.F. & Thomas, J.F. (2000) Reversion of flowering in *Glycine Max* (Fabaceae). American Journal of Botany, 87, 1425–1438 doi:10.2307/2656869

Winiarczyk, K., Marciniec, R. & Tchorzewska, D. (2018) Phenomenon of Floral Reversion In Bolting Garlic (*Allium sativum* L.). Acta Scientiarum Polonorum-Hortorum Cultus, 17, 123–134 doi:10.24326/asphc.2018.2.11

Xi, Y., Fu, C., Ge, Y., Nandakumar, R., Hisano, H., Bouton, J. & Wang, Z. (2009) Agrobacterium-Mediated Transformation of Switchgrass and Inheritance of the Transgenes. BioEnergy Research, 2, 275–283 doi:10.1007/s12155-009-9049-7

Yang, H. (1986) The use of a whole stain-clearing technique for observations on embryo sac, embryo, endosperm and embryoid. Acta Botanica Sinica, 28, 002

Zhao, K., Fan, H., Jiang, X., Zhou, S. (2002) Critical day-length and photoinductive cycles for the induction of flowering in halophyte Suaeda salsa. The Plant Science, 162, 27–31 doi:10.1016/S0168-9452(01)00520-9

Zik, M. & Irish, V.F. (2003) Flower development: initiation, differentiation, and diversification. Annual Review of Cell and Developmental Biology, 19, 119–140 doi:10.1146/annurev.cellbio.19.111301.134635

